# *ramr*: an R package for detection of rare aberrantly methylated regions

**DOI:** 10.1101/2020.12.01.403501

**Authors:** Oleksii Nikolaienko, Per Eystein Lønning, Stian Knappskog

**Author notes:** Corresponding author: Oleksii Nikolaienko, K. G. Jebsen Center for Genome-Directed Cancer Therapy, Department of Clinical Science, University of Bergen, Bergen, Norway.

## Abstract

**Motivation:** With recent advances in the field of epigenetics, the focus is widening from large and frequent disease- or phenotype-related methylation signatures to rare alterations transmitted mitotically or transgenerationally (constitutional epimutations). Merging evidence indicate that such constitutional alterations, albeit occurring at a low mosaic level, may confer risk of disease later in life. Given their inherently low incidence rate and mosaic nature, there is a need for bioinformatic tools specifically designed to analyse such events.

**Results:** We have developed a method (*ramr*) to identify aberrantly methylated DNA regions (AMRs). *ramr* can be applied to methylation data obtained by array or next-generation sequencing techniques to discover AMRs being associated with elevated risk of cancer as well as other diseases. We assessed accuracy and performance metrics of *ramr* and confirmed its applicability for analysis of large public data sets. Using *ramr* we identified aberrantly methylated regions that are known or may potentially be associated with development of colorectal cancer and provided functional annotation of AMRs that arise at early developmental stages.

**Availability and implementation:** The R package is freely available at https://github.com/BBCG/ramr

## Introduction

Epigenetics, normally assessed as gene promoter CpG methylations, plays a pivotal role to many physiological processes throughout life (Fraga et al., 2005). In addition, disturbances in epigenetic function is involved in many pathophysiological processes. Somatic epimutations are frequently seen in cancers (Peltomäki, 2012), and constitutional epimutations (Sloane et al., 2016) have been associated with elevated risk of cancer (Dobrovic and Kristensen, 2009; Evans et al., 2018; Hitchins et al., 2007; Lynch et al., 2015; Prajzendanc et al., 2020) as well as other diseases (Evans et al., 2007; Sloane et al., 2016). Notably, recent findings (Lønning et al., 2018; Lønning et al., 2019) show that even epimutations occurring at a low mosaic level (affecting only a few percent of normal cells) are associated with an elevated cancer risk. While to this end such low-level mosaic methylation has been confirmed for a few genes only (Lønning et al., 2019), the findings are suggestive that similar epimutations occur for several other tumour suppressor genes in respect to different tumour forms as well. Thus, these preliminary results point towards a new and important field of research that may change our understanding of carcinogenesis as well as the origin of several other diseases profoundly.

Low-level mosaic methylation typically affects <10% of the alleles in samples of pooled normal cells (i.e., tissue biopsy or blood sample) and may easily be overlooked by contemporary screening methods. To overcome such problems, it is crucial to develop new, unbiased, exploratory approaches suitable for identification of rare long-range changes in methylation levels, i.e. aberrantly methylated regions (AMRs). Importantly, the application of such approaches is not restricted to detection of high-level hemiallelic epimutations in tumour suppressor genes and their role for cancer development but can also include discovery of moderate mosaic methylation events underlying any disease or condition. Unsupervised tools for AMR identification could be also useful to assess epimutation burden in individuals.

A number of software tools for the analysis of differentially and variably methylated regions have been developed. While some of these tools, like the DMRcate (Peters et al., 2015), bumphunter (Jaffe et al., 2012) or iEVORA (Teschendorff et al., 2016a), can process data from any source, others, like DiffVar (Phipson and Oshlack, 2014), DMRcaller (Catoni et al., 2018) or DMCHMM (Shokoohi et al., 2018), are limited to processing of bisulfite massive parallel sequencing data (e.g. Bismark output) or BeadChip data only. Importantly, all of these tools are meant to compare two sets of samples and some of them proved to be less sensitive (more robust) to outlier values (Teschendorff et al., 2016b). Thus, there is a need for new tools specifically designed to identify the outliers with respect to AMRs in single/few individual samples in a data set.

We here propose and describe a novel unsupervised method, *ramr*, for search of “rare aberrantly methylated regions” in large data sets. By its design, *ramr* is sensitive to biologically relevant (extended over prolonged genomic regions) outliers and is able to find epigenetic aberrations in one or several samples within the data set, making it suitable for discovery of low-frequency and/or mosaic epimutations. Using simulated data, we compared its performance with some existing methods for search of differentially methylated regions (DMR). We also applied *ramr* and other methods to identify and characterize AMRs in public GEO (GSE51032, GSE105018) and TCGA-COAD data sets.

## Methods

### Data sets and data preprocessing

GSE51032: this data set was used to simulate test data, characterize AMRs and find common aberrations with colorectal cancer patients from TCGA-COAD data set. The public GSE51032 data set contains whole blood cells DNA methylation data generated by the Infinium Human Methylation 450 Bead Chip array (Cordero et al., 2015) from 845 participants in the EPIC-Italy cohort (total n=47 746). EPIC is a prospective cohort study designed to investigate the relationship between genetic and environmental factors and the incidence of cancer and other diseases (Riboli et al., 2002). At the time of the most recent follow-up in 2010, 235 of GSE51032 participants had developed incidental breast cancer, 166 incidental colorectal cancer, while n=20 had developed other primary cancers. Blood samples from these 421 patients collected prior to their cancer diagnosis were analysed together with samples from 424 control participants remaining cancer-free. The raw Illumina Infinium HumanMethylation450 BeadChip data files for this data set were obtained from Gene Expression Omnibus (GEO, https://www.ncbi.nlm.nih.gov/geo) and processed (normalized and annotated) with the minfi Bioconductor package (Aryee et al., 2014) using the preprocessQuantile method with outlier thresholding enabled.

The full data set (485 512 CpGs x 845 samples) was first used as a template to create a test data set (see Preparation of test data sets, below). For further analyses (other than generation of test data set), all probes mapping to chromosomes X and Y (using hg19 genome assembly) together with non-specific or polymorphic probes (Chen et al., 2013) were removed prior to identification of aberrantly methylated regions, resulting in a smaller data set with 383 788 CpGs (i.e. 383 788 × 845).

TCGA-COAD: this data set was used to identify AMRs presumably undergoing positive selection during carcinogenesis. The Cancer Genome Atlas (TCGA, https://cancergenome.nih.gov/) processed Illumina Infinium HumanMethylation450 data files for 38 adjacent mucosa and 40 corresponding colon adenocarcinoma samples were obtained from The Genomic Data Commons data sharing platform (https://portal.gdc.cancer.gov/). Probes with beta values missing in more than the half of the samples, non-specific or polymorphic probes or probes mapping to chromosomes X and Y (as described above) were filtered out resulting in a data set with 394 360 CpGs × 78 samples.

GSE105018: this data set was used to gain insight on the potential mechanisms of AMR formation. The Environmental Risk (E-Risk) Longitudinal Twin Study data set contains DNA methylation data obtained by analysing blood samples of 732 complete twin pairs at the age of 18 years (426 monozygotic (MZ) and 306 same-sex dizygotic (DZ) twin pairs) and 194 participants whose co-twin did not have complete data using Infinium Human Methylation 450 Bead Chip array (Hannon et al., 2018). Preprocessed files with normalized beta values for this data set were obtained from GEO (https://www.ncbi.nlm.nih.gov/geo/) and filtered as described above, resulting in a data set with 367 522 CpGs x 1658 samples.

### Preparation of test data sets

For the purpose of serving as a test set, a methylation array data was simulated using the full, unfiltered GSE51032 data set as a template. For each CpG in the GSE51032 data set the parameters of beta distribution were estimated using the ebeta function of the EnvStats R module (Millard, 2013). Using the obtained parameters, 100 random beta values were produced by means of EnvStats::rbeta function. The resulting data set contained 485 512 rows (CpGs) and 100 columns (samples).

To make a list of all potentially methylated regions, CpGs were merged within a window of 1000bp and resulting regions containing at least 10 CpGs per region were kept.

In order to simulate rare methylation events, 2000 regions were randomly selected. Of those, 1000 were uniquely assigned to samples (each of these 1000 regions was assigned to a single sample, 10 regions per sample), while the other 1000 were assigned in a non-unique manner (each of these 1000 regions was assigned to three samples, 10 regions per sample). Thus, every sample in a data set possessed 10 unique and 10 non-unique regions with aberrant methylation.

Next, all CpG beta values corresponding to particular region/sample pair were increased or decreased (depending on overall methylation level of this region) by particular delta (0.025, 0.05, 0.10, 0.25, 0.50). All values below 0.001 were set to 0.001, and all values above 0.999 were set to 0.999. These modified regions thus are ground true positive unique (uGTP) or non-unique (nGTP) regions and were used to assess performance of different methods. Examples of original and modified uGTP/nGTP regions are given in Supplementary Fig. 1.

### *ramr* implementation

Three independent filtering methods for identification of AMRs were implemented.

“IQR”: for every genomic position, median beta value and interquartile range across the sample set were calculated. Then, all data points differing from median by less than a certain (user-defined) number of interquartile ranges were considered non-significant and filtered out. “beta” or “wbeta”: non-weighted or weighted beta distribution, respectively, was fit to data for every genomic position, and probability value was calculated for every data point. For weighted parameter estimation, individual values were split in bins, and their assigned weights directly correlated with the number of values in the same bin, and inversely – with the absolute difference from the median value, resulting in lower p-values for outliers. Then, data points with probability values above a certain threshold were considered non-significant and filtered out.

After filtering, significant per-sample data points obtained by selected filtering method were then merged by genomic position within a particular window, and aggregate p-value was calculated as geometric mean of p-values for individual significant data points comprising genomic region. Additional filtering was then applied to the list of aberrantly methylated genomic regions (by minimum number of merged CpGs, minimum average difference from median beta value, etc).

### Method comparison

The test data was analysed by *ramr* with the following parameters: filtering by IQR (method=”IQR”) or non-weighted (method=”beta”) or weighted (method=”wbeta”) beta distribution fitting, IQR cutoff or q-value cutoff as specified (range of 2 to 9, or 1e-02 to 1e-09, respectively), merging CpGs within 1000bp (merge.window=1000), selecting AMRs with at least 5 significantly different beta values (min.cpgs=5), using 7 parallel threads (cores=7).

We compared *ramr* to the following methods widely employed for differential methylation analysis: dmpFinder (R package minfi v1.30.0 with qCutoff as specified and other default parameters), champ.DMP (R package ChAMP v2.14.0 with adjPVal as specified and other default parameters (Morris et al., 2013)), champ.DMR (R package ChAMP v2.14.0, with the following parameters: adjPvalProbe as specified, method=“ProbeLasso” minDmrSep=1000, meanLassoRadius=1000), lmFit (R package limma v3.39.19 with the default parameters (Ritchie et al., 2015)) followed by comb-p (python module v0.50.3, with the following parameters: seed as specified, dist=1000 (Pedersen et al., 2012)). As these methods require two classes/categories for comparison, every sample from the test data set was tested against all the other samples. For dmpFinder and champ.DMP, differentially methylated CpGs detected were merged and filtered as described for *ramr* above. Probability cutoff value for all of the existing methods was in range of 1e-02 to 1e-09.

True positive unique (uTP) or non-unique (nTP) region is defined as region which overlaps by at least 1 bp with any of ground true positive unique (uGTP) or non-unique (nGTP) regions correspondingly. False positive (tFP) region is defined as region not overlapping with any of uGTP or nGTP regions.

The following metrics were calculated:

1) Precision (Positive Predictive Value):

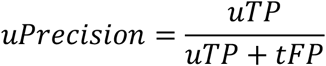

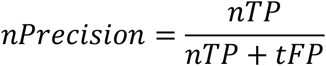
2) Recall (True Positive Rate):

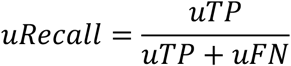

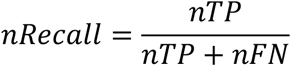

where uFN and nFN are the number of unique or non-unique false negative regions, respectively.
3) Matthews correlation coefficient (MCC):

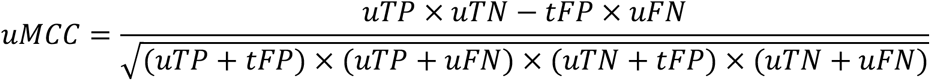

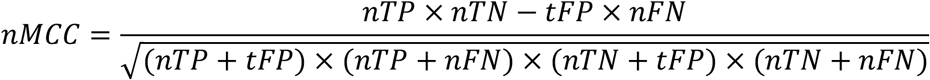

where uTN and nTN are the number of unique or non-unique true negative regions, respectively.
4) F1 score

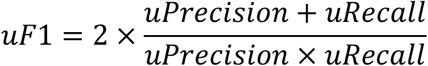

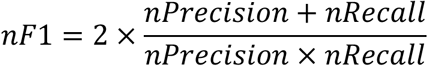
5) AuPR uAuPR (for unique regions) and nAuPR (for non-unique regions) values were calculated using R package PRROC (v1.3.1) using precision and recall metrics as calculated above.
6) Time The elapsed time measured in seconds for AMR search using every algorithm was recorded on a HP OptiPlex 7050 PC with 64 GB 2400 MHz, DDR4 RAM, 8-core Intel® Core™ i7-7700 (3.60GHz) CPU and the Ubuntu 18.04.4 LTS Operating System. Due to relatively low performance of some methods and multiple test scenarios, parallel computing on 7 cores was used when comparing computing times for different algorithms. As comb-p uses maximum of four threads by default, single threading was forced to obtain results comparable to ChAMP and minfi. Then, multiprocessing of all methods except *ramr* was achieved by running 7 independent processes at a time. Method performance in a single-process mode was also assessed for several test scenarios and was found to be consistent with multi-process estimates.

### Identification and characterization of aberrantly methylated regions

Pre-filtered data sets (see above) GSE51032 (383 788 CpGs, 845 samples), TCGA-COAD (394 360 CpGs, 38 adjacent mucosa samples), and GSE105018 (367 522 CpGs, 1658 samples) were analysed using *ramr* with the following parameters: method=”beta”, qval.cutoff=1e-3, min.cpgs=7, merge.window=1000.

### Region annotation and enrichment analysis

Genomic regions were annotated using the R package annotatr v1.10.0 (Cavalcante and Sartor, 2017). Chromatin marks overrepresented in aberrantly methylated regions were assessed by locus overlap analysis for enrichment of genomic ranges (R package LOLA v1.14.0 (Sheffield and Bock, 2015)) using a provided extended data set containing 1110 BED files from the Roadmap Epigenomics Project. Briefly, every given set of AMRs was tested for enrichment in chromatin marks using function runLOLA with redefineUserSets parameter set to TRUE. Significant hits (qValue<1e-3) were grouped by chromatin mark-specific antibody and counted. A set of genomic regions which was used as a reference set for annotation and enrichment analysis (“universe”) was obtained by merging genomic positions of GSE51032 data set probes with the following parameters: min.cpgs=7, merge.window=1000.

For enrichment analysis using chromatin marks in developing mouse embryo, both the specified AMRs and the “universe” set were lifted over to mm9 mouse assembly using R package liftOver v1.8.0 (https://www.bioconductor.org/help/workflows/liftOver/). A data set containing mouse genomic regions carrying H3K4me3, H3K9me3 or H3K27me3 for various developmental stages was obtained from GEO (accession number GSE98149; https://www.ncbi.nlm.nih.gov/geo).

## Results and discussion

Merging evidence indicates that constitutional mosaic epimutations arising in early embryonic life may be a risk factor for certain cancer forms (Lønning et al., 2019) as well as other diseases (Evans et al., 2007) later in life. However, such epimutations are rare, and may be difficult to identify comparing pooled subsets of cases and controls. In order to detect the genomic regions aberrantly methylated in a small subset of samples, we developed a fast method for within-the-class differential methylation analysis, omitting the need of splitting samples in subsets for comparison.

### Implementation and evaluation of the method

Assuming beta distribution of methylation values, for every sample at every given genomic position the method estimates distribution parameters and calculates either p-value, or deviation (xiqr) from the median value normalized by interquartile range (IQR). Filtering by p-values or xiqr is applied, and significant genomic positions that remain after filtering are merged into regions using floating window of a provided length. Thereafter, post-filtering is performed to select for regions bearing no less than a specified number of significant genomic positions, followed by a calculation of per-region p-values (see Methods section for details). The implemented method (“rare aberrantly methylated regions”; *ramr*) accepts GRanges object, containing beta values as metadata columns (samples), and returns GRanges object with all the AMRs identified in any of the samples. By its design, the method is not constrained by the source of input data and is suitable for analysis of data obtained by methylation profiling using both array and next-generation sequencing of non-methylation-enriched samples.

In order to evaluate sensitivity and specificity of our approach versus existing methods, we simulated 450k array data using GSE51032 as a template (see Methods section for details). All three *ramr* filtering methods (“IQR”, “beta” and “wbeta”) were applied to find artificially introduced AMRs in the simulated data set. The performance was compared to four other available methods (champ.DMP, champ.DMR, lmFit + comb.p and dmpFinder). Accuracy metrics and computing times were also compared using the simulated data set. To select the best performing method we used Matthews correlation coefficient (MCC) as the most reliable metric for classifications of imbalanced sets (Chicco and Jurman, 2020).

The top results from the test runs are summarized in Table 1 with further detail in Supplementary Table 1. The results indicate that *ramr* and comb-p performed similarly and consistently better than the other methods for differential methylation analysis across all simulation scenarios (as previously revealed by (Mallik et al., 2019)). comb-p gave better results than *ramr* when moderate changes in beta values were introduced (delta=0.05-0.25 for unique or 0.1-0.25 for non-unique regions), while *ramr* outperformed comb-p for very small and large methylation aberrations. Computing times for *ramr* were 60x-240x lower than for comb-p (Fig. 1, Table 1), making it more suitable for analysis of large data sets with only a marginal loss in balanced accuracy compared to comb-p. Of note, comb-p, in contrast to other methods, has no parameter to filter the regions based on number of significant genomic positions, which may explain higher number of true positives and false negatives obtained.

**Table 1.** Top Matthews correlation coefficient (MCC) values for unique (uMCC) and non-unique (nMCC) AMR identification are given in bold, corresponding cutoff values – in parentheses.

|  | delta |  |  |  |  |  |  |  |  |  |
| --- | --- | --- | --- | --- | --- | --- | --- | --- | --- | --- |
|  | 0.025 |  | 0.050 |  | 0.100 |  | 0.250 |  | 0.500 |  |
| method | uMCC | nMCC | uMCC | nMCC | uMCC | nMCC | uMCC | nMCC | uMCC | nMCC |
| dmpFinder | NA | NA | 0.0707 | 0.0182 | 0.7569 | 0.4526 | 0.9803 | 0.9633 | 0.9884 | 0.9861 |
| champ.DMP | NA | NA | 0.0632 | NA | 0.7582 | 0.4548 | 0.9803 | 0.9661 | 0.9884 | 0.9864 |
| champ.DMR | NA | NA | NA | NA | 0.4266 | 0.2235 | 0.7402 | 0.6533 | 0.7674 | 0.7214 |
| lmFit + comb.p | 0.1336 | 0.1608 | **0.7663**<br>(1e-06) | 0.7514 | **0.9711**<br>(1e-08) | **0.9630**<br>(1e-07) | **0.9950**<br>(1e-09) | **0.9916**<br>(1e-07) | 0.9990 | 0.9993 |
| *ramr* (iqr) | **0.1980**<br>(2) | 0.1732 | 0.7348 | 0.7322 | 0.9644 | 0.9584 | 0.9879 | 0.9889 | 0.9985 | 0.9968 |
| *ramr* (beta) | 0.1488 | 0.1576 | 0.7351 | 0.7092 | 0.9444 | 0.9513 | 0.9889 | 0.9849 | **1.0000**<br>(1e-03) | 0.9983 |
| *ramr* (wbeta) | 0.1573 | **0.2107**<br>(1e-03) | 0.7246 | **0.7741**<br>(1e-03) | 0.9560 | 0.9497 | 0.9894 | 0.9896 | **1.0000**<br>(1e-04) | **0.9997**<br>(1e-04) |

**Fig. 1.**
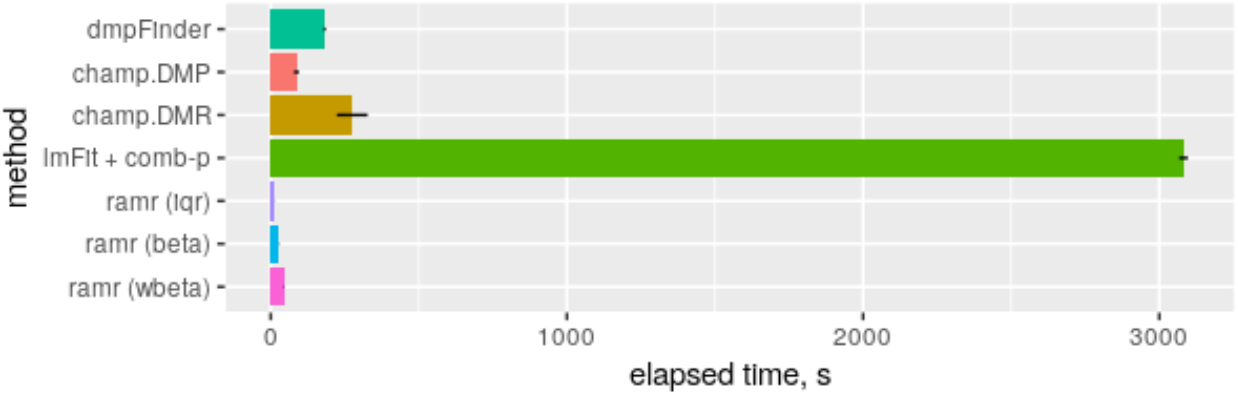
Performance of different methods. Computational time was measured as described in Methods section.

The three different filtering techniques in *ramr* vary in their precision/recall metrics and were implemented in parallel in order to provide high degree of analysis flexibility. As confirmed by performance evaluation using the simulated data set, IQR filtering is the fastest and the most stringent but, at the same time, the least sensitive (lower number of TP and FP) among the *ramr* filtering methods. In contrast, fitting weighted beta distribution increases computational time as well as the number of true positive and false negatives, while fitting non-weighted beta distribution provide a balance between speed and accuracy. In addition, performance metrics vary for unique and non-unique AMRs, thus the best parameters are to be estimated for any particular analysis case.

### Characterization of aberrant methylation events in the EPIC-Italy sample set

Aiming to characterize real AMRs, we applied *ramr* to several publicly available methylation data sets. As methylation variation of individual probes may be a result of technical errors or nucleotide polymorphism, we decreased p-value threshold to 1e-3 and limited our analysis to genomic regions containing at least 7 aberrantly methylated CpGs, which is also thought to enhance the biological relevance of search hits.

The search for AMR in the EPIC-Italy GSE51032 data set resulted in 3582 AMRs across the 845 samples, 2888 of them being hypermethylation- and 694 hypomethylation events. The AMRs were unevenly distributed across chromosomes, occurring at high frequencies on chromosomes 6 and 11 versus particularly low frequency on chromosome 9 (Fig. 2A).

**Fig. 2.**
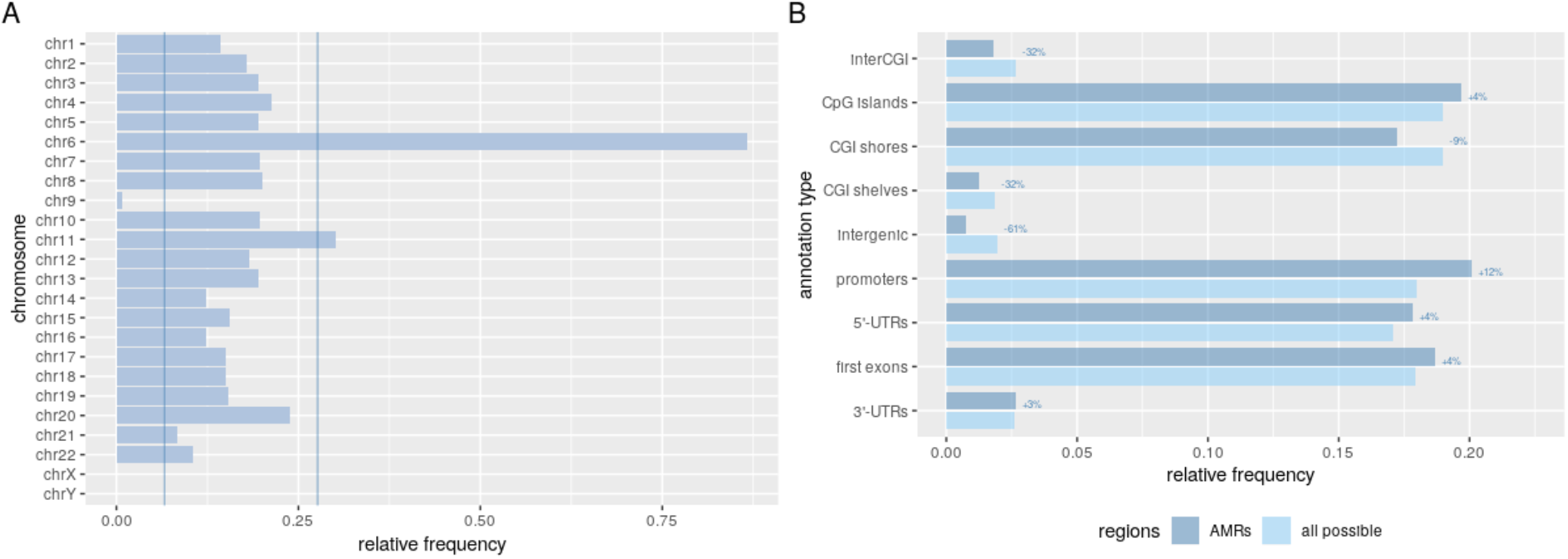
(A) Distribution of AMRs across chromosomes. Number of identified AMRs was normalized by the possible number of regions per each chromosome. Vertical lines mark frequency values equal to Q_1_– 1.5*IQR and Q_3_+1.5*IQR; (B) AMR distribution across various genomic regions. Structural annotations of AMRs or all possible regions were summarized and normalized to their total number. Labels represent percent change per each annotation category.

To further characterize the identified AMRs, we annotated them by their positions relative to known genomic elements. Compared to all possible genomic regions, represented within the GSE51032 data set, AMRs were detected more frequently at core CpG islands and 5’-UTRs (in general associated with gene promoter regions) compared to distant CpG island elements (shores and shelves) or other intergenic regions (Fig. 2B). Per-sample number of AMRs was in a wide range from 0 to 602 with a mean value of 4 and median value of 1 AMR per sample (Fig. 3). Assuming normal distribution of a number of per-sample aberrant methylation events, we classified samples into low-AMR and high-AMR groups using a simple outlier detection rule (threshold=Q_3_+1.5*IQR). High-AMR samples (n=44) carried most of the AMRs identified (n=2328).

**Fig. 3.**
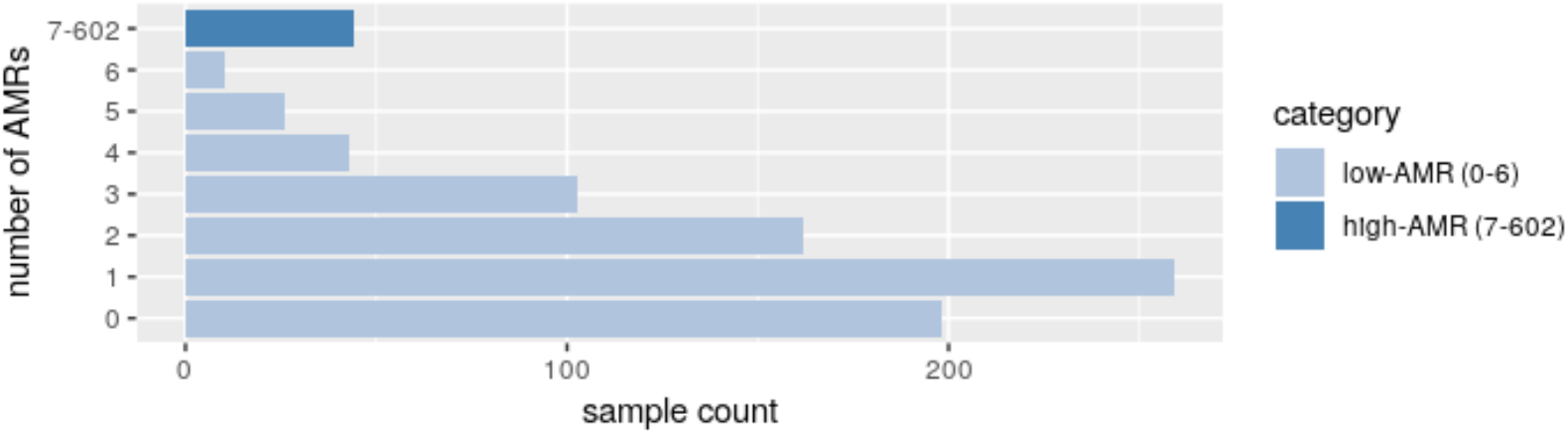
Sample count distribution in low- and high-AMR sample groups.

Using information on age of participants (patients and controls together), we confirmed an age-related increase in the number of AMRs (previously established in (Fraga et al., 2005)), but only in the subset of low-AMR individuals (p=0.000762, vs p=0.336 for all participants), indicating that very high number of AMRs in some of the samples may not relate to aging, but could result from deregulation of the epigenetic machinery in hematopoietic cells. Of note, none of the participants from the GSE51032 data set developed malignant neoplasms of lymphoid, hematopoietic or related tissue, known to be often associated with methylation alterations (Vosberg et al., 2019), at the time of the most recent follow-up. Further subdividing AMRs from low-AMR individuals according to their genomic annotations revealed that the long-range age-related changes accumulate mostly within CpG island shores (Bonferroni-corrected p=0.00246), while other genomic elements did not show significant correlation between age and number of AMRs (similar correlation was previously reported in (Slieker et al., 2018), reviewed in (Unnikrishnan et al., 2019)).

We further assessed enrichment of AMRs in epigenetic marks using histone modification patterns of 111 reference human epigenomes from Roadmap Epigenomics Human Epigenome Atlas (Kundaje et al., 2015). The full set of AMRs, as well as AMRs subsets belonging to high- or low-AMR samples, showed enrichment in various chromatin modifications which mark active or repressed chromatin (Fig. 4). To assess variability between individual samples we performed similar analysis for the 10 most AMR-rich samples from the high-AMR subset. Differences in their enrichment patterns confirm the existence of multiple aberration types that may cause AMR accumulation in individuals – such as overexpression of DNMT enzymes (Zhang et al., 2018) or mutations in their DNA-recognizing domain (Sendžikaitė et al., 2019).

**Fig. 4.**
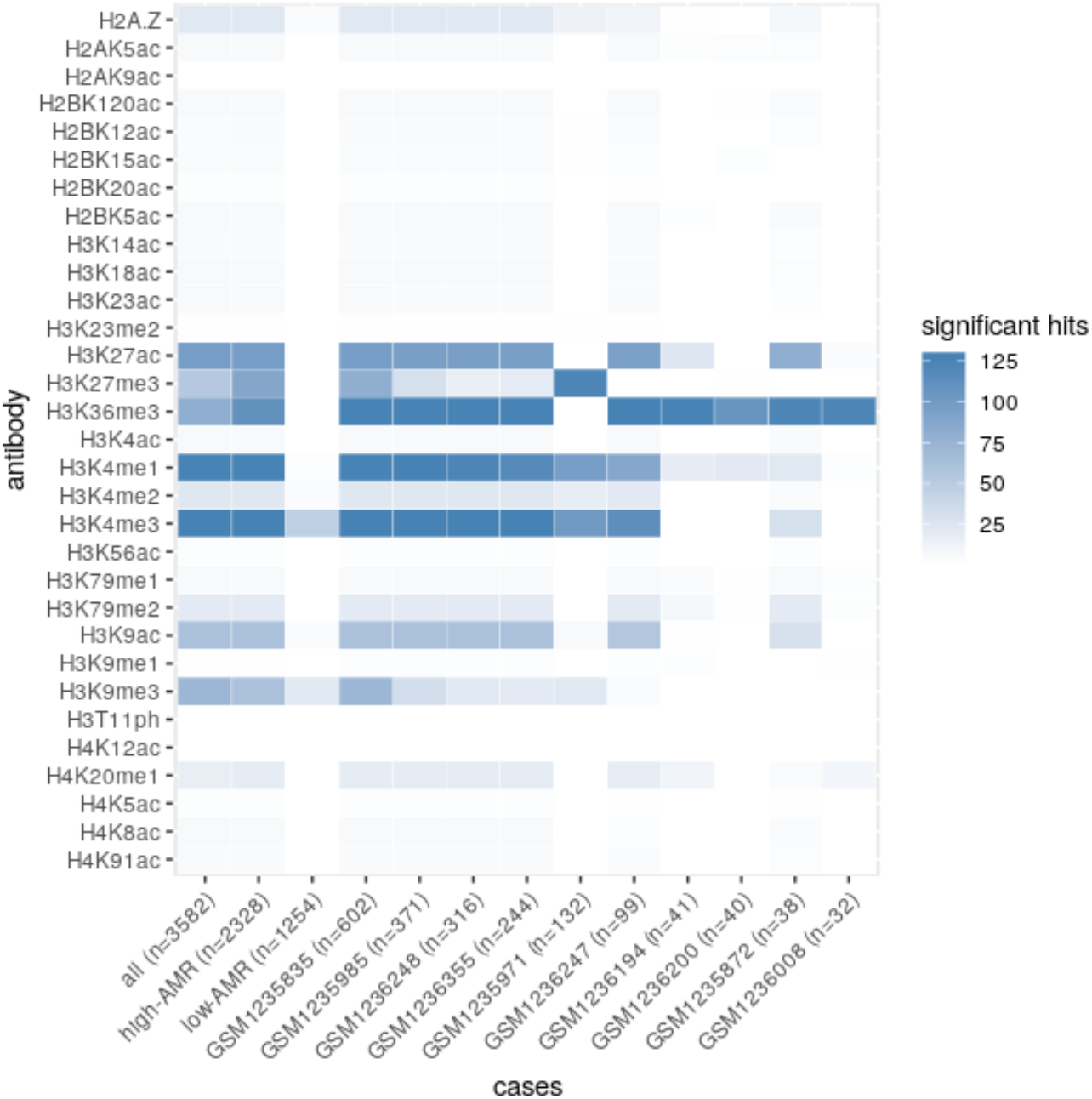
Heatmap plot of AMR enrichment in chromatin modifications. AMRs belonging to particular sample groups or individual high-AMR samples were checked for enrichment in known chromatin modifications. Heat map shows summarized number of significant hits per sample or sample group.

### Potential cancer-inducing AMRs

Local epigenetic alterations are known to accumulate and clonally expand in normal mitotic tissues (Bian et al., 2002; Graham et al., 2011; Li et al., 2016). Together with genetic alterations they underlie field cancerization phenomenon best described in gastrointestinal cancers (Baba et al., 2016). Numerous frequent events were already reported to be associated with carcinogenesis or risk of cancer (Sakai, 2014; Takeshima and Ushijima, 2019). In order to predict yet unknown, potential cancer-inducing aberrant methylation events, we performed a search of AMRs in methylation profiles of a subset of the TCGA-COAD data set containing adjacent normal mucosa samples from patients with colon cancer (n=38). As cancer-inducing aberrant methylation events are expected to be positively selected during carcinogenesis, we post-filtered the set of AMRs using the following criteria: 1) AMR methylation levels of corresponding tumour and adjacent mucosa samples must deviate in the same direction, and 2) absolute difference between AMR methylation levels of corresponding tumour sample and adjacent mucosa sample must be greater than 0.2. To find out which of the selected regions may exist as cancer-predisposing epimutations, we checked if aberrant methylation in those genomic regions was detected in a subset of low-AMR individuals from GSE51032 data set that have developed colorectal cancer. Four such regions have been found (none of them coappeared in the same sample): chr3:37033791-37035399 (*EPM2AIP1*, *MLH1*), chr12:133463694-133464933 (*RP11*-*46H11.12, CHFR*), chr6:31783029-31783545 (*HSPA1L*, *HSPA1A*), chr19:28284491-28285308 (*LINC00662*, *CTC-459F4.3*, *LLNLF-65H9.1*). Interestingly, constitutional epimutation in one of these regions which belong to *MLH1* gene has been established as rare cause of Lynch syndrome (Lynch et al., 2015), the *CHFR* genomic region has been found to be hypermethylated in colorectal cancer tissue (Sun et al., 2017), *HSPA1A* – in ovarian and bladder cancers (Ban et al., 2019; De Andrade et al., 2019), while *LINC00662* has been shown to promote tumourigenesis in colorectal cancer (Wang et al., 2020). To the best of our knowledge, potential risk for colorectal cancer related to constitutional mosaic epimutations in any of these genes have not been formally assessed. Of note, we did not identify normal tissue aberrations in other known genes in which methylation is known to be frequent in development of colorectal cancer, such as *MGMT* (Menigatti et al., 2009) and *MSH2* (Kang et al., 2015), likely due to the limited number of samples analysed (n=38). Further studies are needed in order to detect other rare events or investigate their potential effect on cancer risk.

### Aberrant methylation events in twins

The emergence of aberrant methylation events may be a result of genetic and/or environmental factors and may potentially occur during various stages of development. Therefore, methylation outlier regions may be present throughout all normal tissues (germline epimutations), in some specific organs, or show mosaic distribution (later developmental or clonal expansion events) (Fraga et al., 2005; Takeshima and Ushijima, 2019). In order to gain more insight into the potential mechanisms of formation of AMRs, we assessed methylation aberrations in a large set of twins from the Environmental Risk (E-Risk) Longitudinal Twin Study (GSE105018). Of all the twin pairs in the data set (426 monozygotic (MZ) and 306 dizygotic (DZ) pairs), 238 MZ and 142 DZ twin pairs consisted of individuals both having at least one AMR each (the mean number of AMRs per individual was 1.60 and 1.76 for MZ and DZ twins, respectively). One hundred and seventy-six MZ and 46 DZ pairs had at least one AMR overlapping between the two individuals in the pair (the mean number of overlapping AMRs between twins were 1.11 and 0.35 for MZ and DZ twin pairs, respectively). Nearly two-fold difference between the relative frequencies of pairs with overlapping AMRs (176/238=0.74 and 46/142=0.32, for MZ and DZ twins, respectively) suggests that the emergence of many overlapping AMRs is possibly triggered by genetic components. At the same time, there is also a substantial number of non-overlapping AMRs in both MZ and DZ twins, implying frequent stochastic events.

For all twin pairs having at least one AMR in common, the mean number of common AMRs was 2.54 and 2.51 for MZ and DZ individuals, respectively. In comparison, the mean number of AMRs overlapping between individuals from different twin pairs was 0.044 and 0.039 for MZ and DZ subjects, respectively. Interestingly, methylation profiles in AMRs were similar within twin pairs sharing the AMRs: using only the AMRs genomic coordinates and their methylation profiles we were able to correctly identify corresponding twins for 234 individuals from 123 MZ and 16 DZ twin pairs. There were no gender-specific differences with respect to the above mentioned AMR frequencies or overlap between individuals.

Additionally, according to enrichment analysis, overlapping AMRs – which are thought to emerge during early development – showed significant and exclusive enrichment in H3K9me3 marks in human fetal tissues (adjusted p=1.29e-5 or higher). The same overlapping AMRs being lifted over to mouse assembly showed even stronger enrichment in H3K9me3 marks in 7.5-days mouse embryo (Bonferroni-adjusted p=4.60e-25). Interestingly, di- and trimethylation of H3K9 is known to protect maternal 5-methylcytosine from oxidation and subsequent demethylation in the zygotes (Wang et al., 2018; Zeng et al., 2019), hinting towards potential involvement of aberrant H3K9 methylation in AMR emergence after fertilization. Taken together, these findings indicate that genetic and/or early environmental influence is dominating the generation of AMRs in individuals, and that aberrantly methylated genomic regions often bear specific epigenetic patterns.

## Conclusion

Involvement of epigenetic alterations in the development of various diseases have been previously demonstrated and is being confirmed in increasingly larger-scale epigenetic studies (Wong et al., 2020). It is also predicted that epigenetic alterations in a higher than currently anticipated number of cancer-predisposing genes might affect cancer risk (Widschwendter et al., 2018). Given the rapid evolution and cost reduction of next-generation sequencing (NGS), including its widespread use in epigenetics, an increasing number and scale of studies in this area are expected. Consequently, bioinformatic tools allowing versatile analysis of resulting data sets will be of critical importance. We believe that our unbiased approach for rare AMR discovery, which is applicable to both array and NGS data, will help with generation of hypotheses and aid in discovery of more disease risk-related epigenetic aberrations.

## Acknowledgements

The authors would like to thank Professor Jan Bulla for valuable early discussions on the topic and computing efficiency of the algorithm.

## Funding

The present work was performed in the Mohn Cancer Research Laboratory and was supported by funding from the K.G. Jebsen foundation, the Norwegian Research Council, The Norwegian Cancer Society and the Norwegian Health Region West.

## Conflict of Interest

none declared.

## Data availability

The data underlying this article are freely available at DataverseNO research data repository, https://doi.org/10.18710/ED8HSD.

---

Supplementary Table 1. Elapsed time in seconds, GTP, TP, FP, GTN, FN, TN, Precision, Recall, MCC, F1 and AuPR values for unique and non-unique AMRs for the different AMR detection methods across multiple cutoff values based on simulation data sets.

Supplementary Fig. 1. Methylation profiles of eight random genomic regions from the GSE51032 template data set (“GSE51032”) together with the methylation profiles of the same regions in each of five simulated test data sets obtained by changing beta values within those regions for a subset of samples (highlighted) by a particular delta (“delta=0.025”, “0.050”, “0.100”, “0.250” or “0.500”).

## Notes

https://github.com/BBCG/ramr

https://doi.org/10.18710/ED8HSD

### Competing Interest Statement

The authors have declared no competing interest.

