## Supplementary figures and images for "*ramr*: an R package for detection of rare aberrantly methylated regions"

### Supplemental Fig. 1

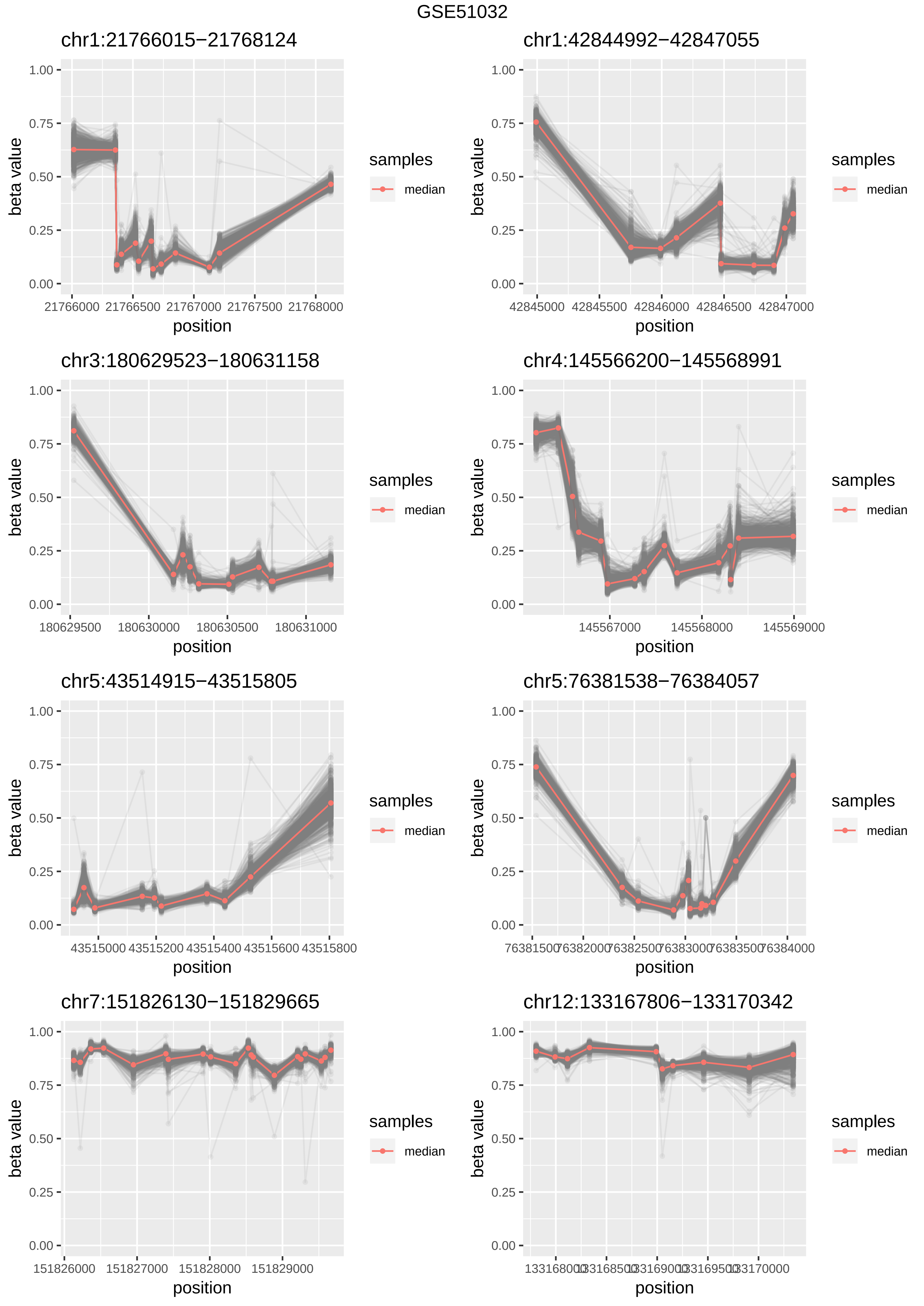

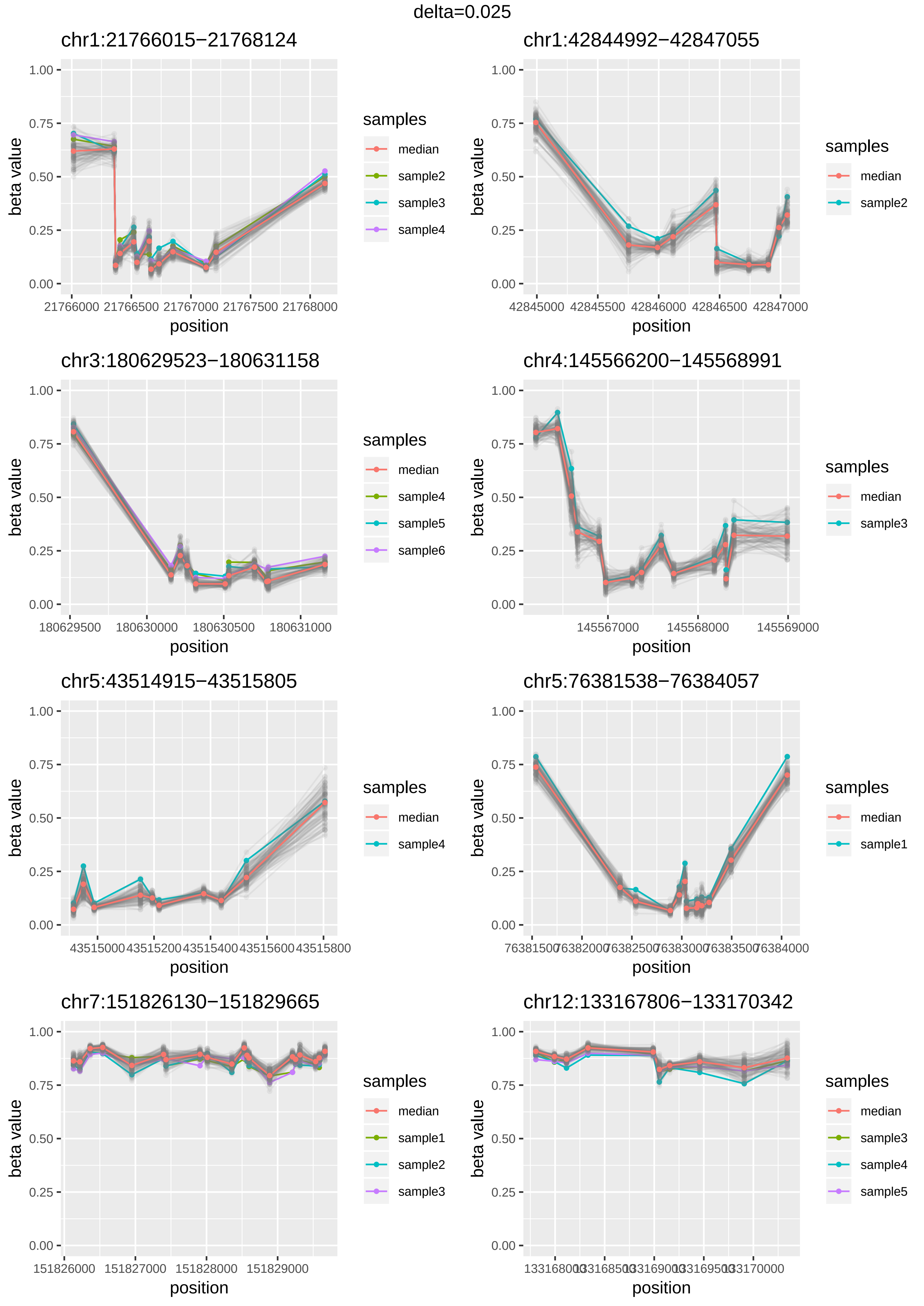

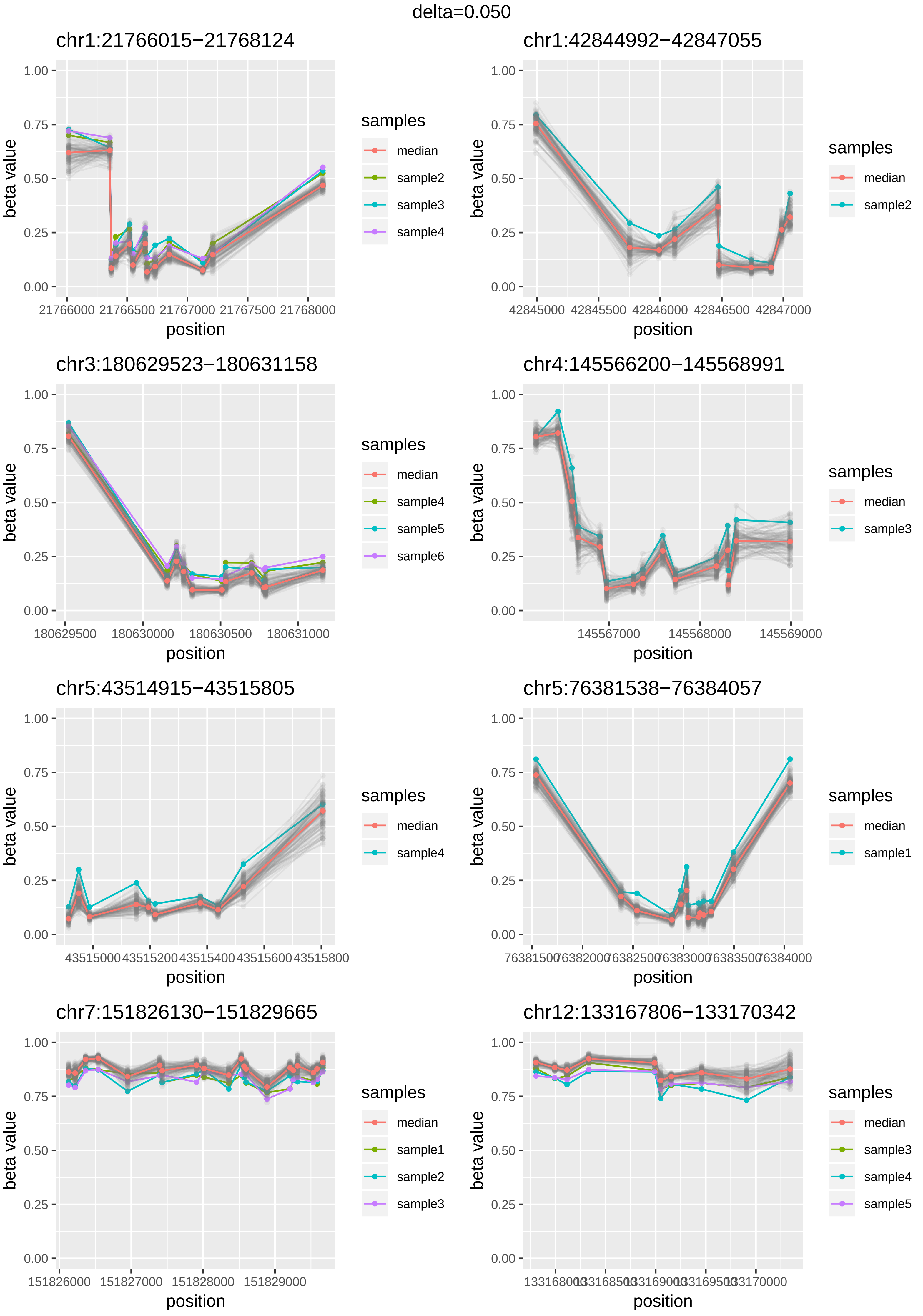

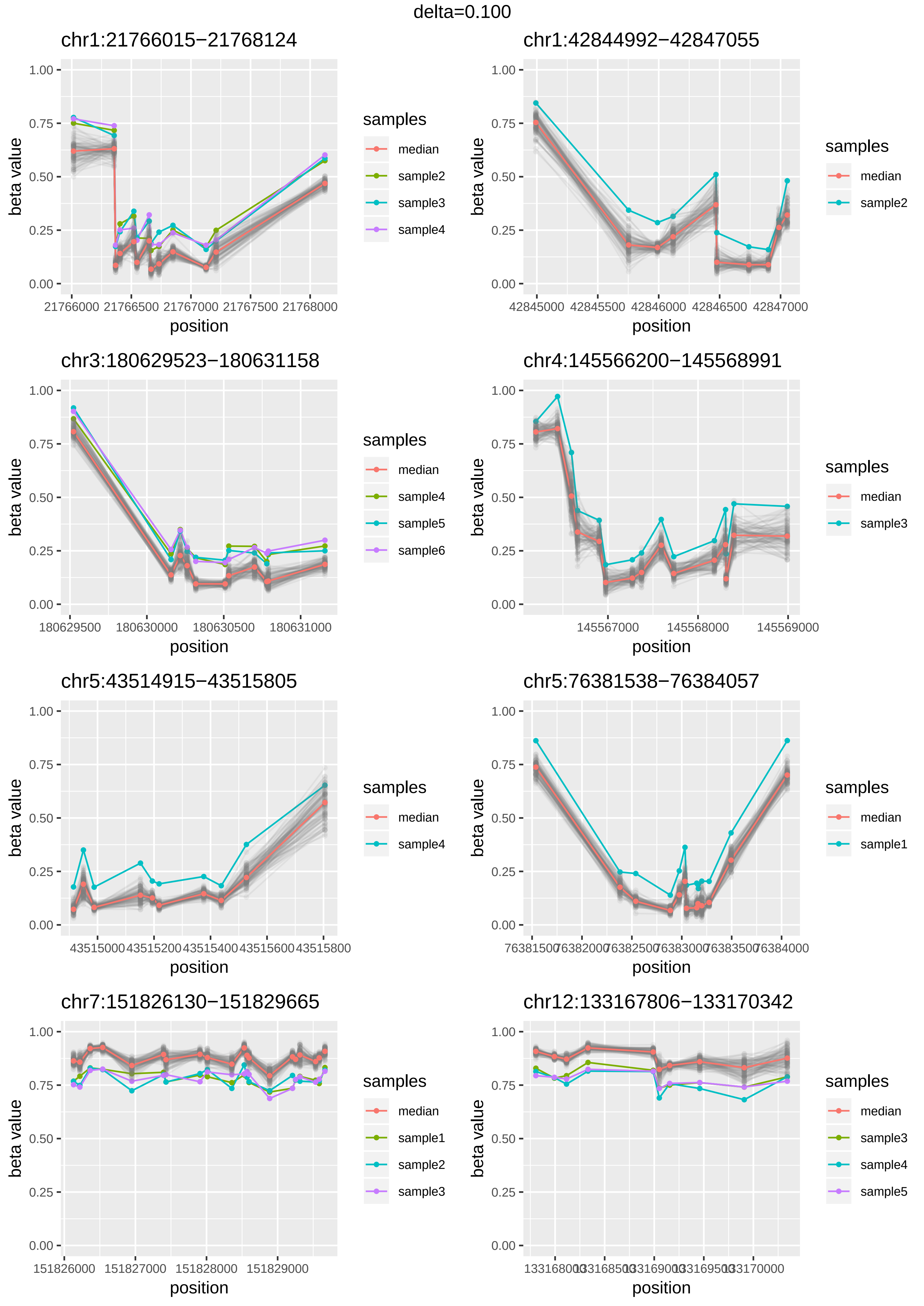

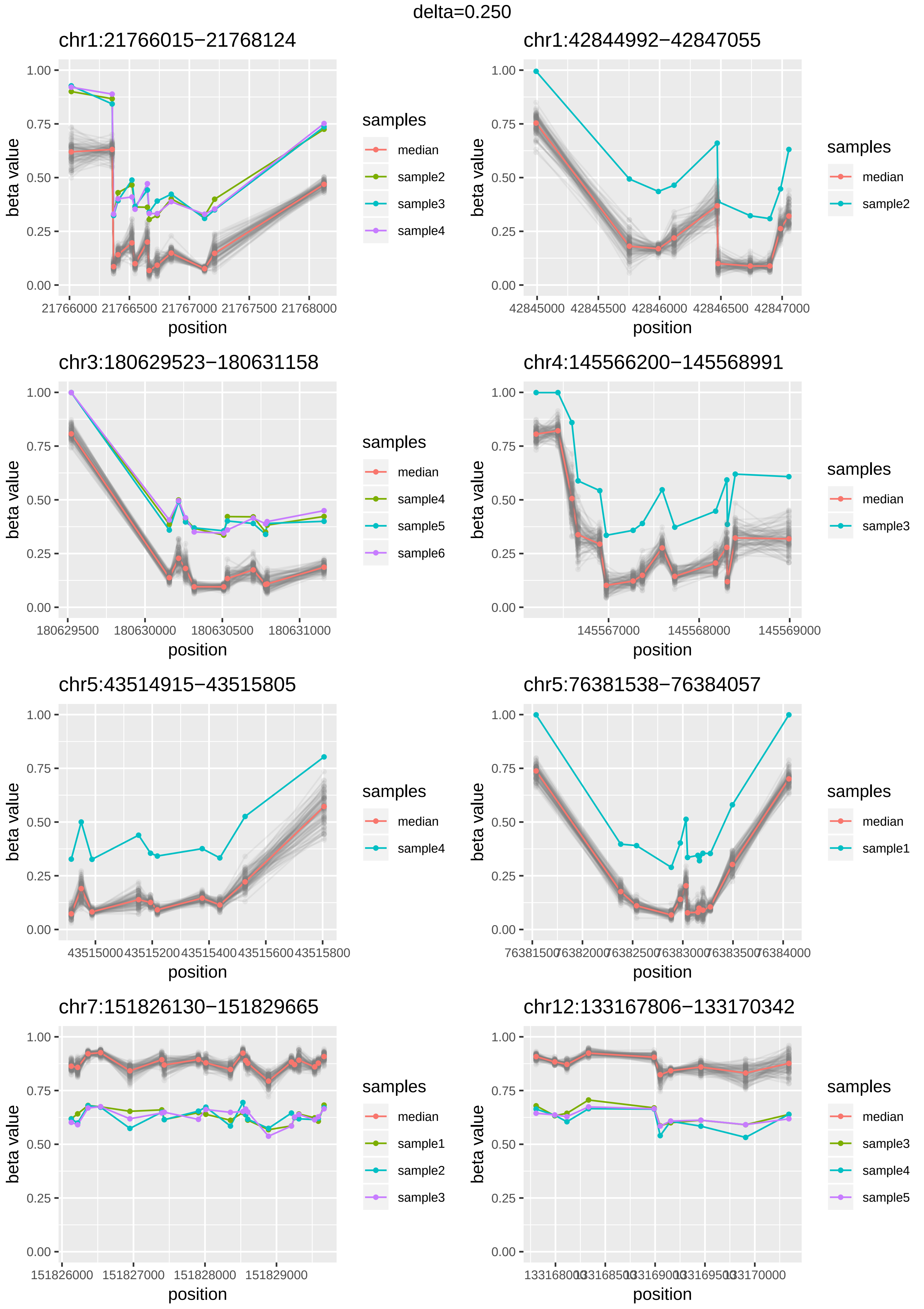

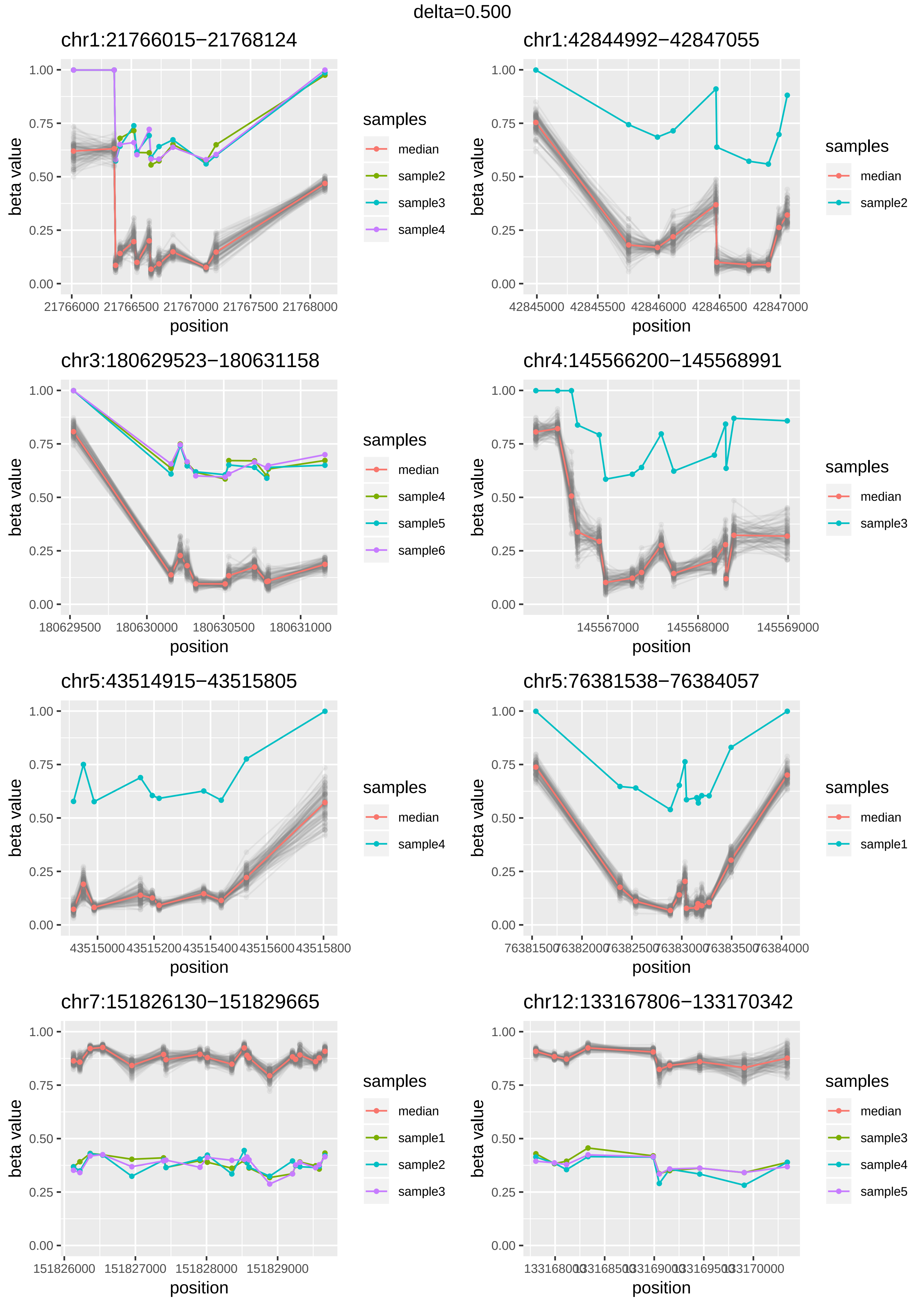
